# Targeted extracellular degradation of LRP8 promotes ferroptosis in cancer cells

**DOI:** 10.64898/2026.05.16.725645

**Authors:** Fangzhu Zhao, Alex Inague, Trenton M. Peters-Clarke, Yifei Chen, Snehal D. Ganjave, Yun Zhang, Kun Miao, Zi Yao, Yan Wu, Madison K.C. Seto, Kevin K. Leung, James A. Olzmann, James A. Wells

## Abstract

Tumor reliance on antioxidant defenses creates a vulnerability to ferroptosis, yet strategies to therapeutically disable these systems remain limited. Here, we identify targeted degradation of the selenium uptake receptor LRP8 as an effective approach to decrease the abundance of the ferroptosis-protective enzyme glutathione peroxidase 4 (GPX4). Using bispecific cytokine receptor–targeting chimeras (KineTACs) that couple LRP8 to cytokine receptor internalization pathways, we selectively direct LRP8 to the lysosome for degradation. LRP8 degradation reduces the abundance of several selenoproteins, including GPX4, lowering the cellular threshold for lipid peroxidation and sensitizing cancer cells to ferroptosis. These findings establish receptor-mediated selenium uptake as a critical, targetable node in ferroptosis resistance and demonstrate that extracellular protein degradation can be leveraged to reprogram intracellular translational dependencies in cancer cells. More broadly, this work provides a framework for exploiting nutrient acquisition pathways to overcome therapy resistance.

## Introduction

Ferroptosis, an iron-dependent form of regulated cell death characterized by the catastrophic accumulation of oxidatively damaged lipids (i.e., lipid peroxides), has emerged as a promising therapeutic target for treatment-resistant cancers(1–3). Several tumor types exhibit heightened sensitivity to ferroptosis induction, such as clear cell renal cell carcinoma(4), MYCN-amplified neuroblastoma(5–8), drug-tolerant persister cells(9–11), and metastasizing cancer cells(12). Ferroptosis induction not only eliminates treatment-refractory cancer populations but may also synergize with existing modalities, such as radiotherapy, immunotherapy, and targeted inhibitors.

Many cancers possess defense systems that promote ferroptosis resistance. The primary mechanism is mediated by the glutathione peroxidase 4 (GPX4)(13, 14), which directly reduces lipid hydroperoxides to inert lipid alcohols. Complementary radical-trapping antioxidant systems, such as the ferroptosis suppressor protein 1 (FSP1)–coenzyme Q10 (CoQ10) pathway(15, 16) and the GTP cyclohydrolase 1 (GCH1)–tetrahydrobiopterin (BH4) axis(17, 18), further act to prevent the propagation of lipid peroxidation. The catalytic activity of GPX4 depends on a selenocysteine residue in its active site, which is incorporated via a specialized translation process that recodes UGA stop codons as selenocysteine insertion sites(19, 20). Adequate selenium supply is therefore essential for sufficient GPX4 expression and activity. Selenium is delivered either as inorganic selenite or through selenoprotein P(21) (SEPP1, also known as SELENOP, which naturally contains 10 selenocysteines) endocytic uptake mediated via its receptors, such as the low-density lipoprotein receptor-related protein 8 (LRP8), and subsequent lysosomal breakdown(19, 22). Recent work indicates that LRP8 is important for tumor growth and ferroptosis resistance, including MYCN-amplified neuroblastoma(22, 23) and triple-negative breast cancer(22–24), acting to provide rapidly growing cancer cells with sufficient selenium pools for GPX4 translation. Loss of LRP8 results in ribosome stalling at the selenocysteine codon of GPX4, ultimately leading to ribosome collisions, premature translation termination, and reduced GPX4 abundance(24). Functionally, the reduction in GPX4 translation sensitizes cells to ferroptosis. Remarkably, in MYCN-amplified neuroblastoma, LRP8 loss is sufficient to trigger ferroptosis both in culture and in orthotopic xenograft models(23), underscoring its potential as a therapeutic target.

Recently advances in extracellular targeted protein degradation (eTPD) have exploited natural endolysosomal and/or proteasomal pathways to eliminate both membrane-bound and soluble proteins(25–30). These bifunctional antibodies harness diverse cell-surface recycling and degradation mechanisms to selectively remove extracellular targets, thereby expanding the druggable proteome to include plasma membrane proteins essential for tumor survival. Building on this concept, we developed bispecific antibody degraders that induce the selective degradation of the SEPP1 receptor, LRP8. By disrupting the SEPP1–LRP8 axis using the cytokine receptor-targeting chimeras (KineTACs)(27), these degraders reduce GPX4 levels, thereby sensitizing cancer cells to ferroptosis. This strategy establishes a tractable framework for therapeutic targeting of ferroptosis resistance and highlights the potential of antibody-based protein degradation to capitalize upon key vulnerabilities in cancer.

## Results

### Engineering antibodies against LRP8

It has been reported that SEPP1 binds to the β-propeller domain of LRP8(31). To target this region, we selected antibodies against the EGF-like domain of LRP8 (LRP8β), which encompasses the β-propeller domain (**Fig. 1A**). We expressed biotinylated LRP8β using a tobacco etch virus (TEV)-Fc-Avitag construct to enable phage clearance via TEV protease cleavage. Phage-displayed Fab libraries were subjected to iterative rounds of selection against immobilized LRP8 antigen, with preclearing steps to remove nonspecific and Fc-binding clones. Enriched binders were recovered after each round and amplified under increasingly stringent conditions to isolate high-affinity, antigen-specific antibodies(32).

**Fig. 1.**
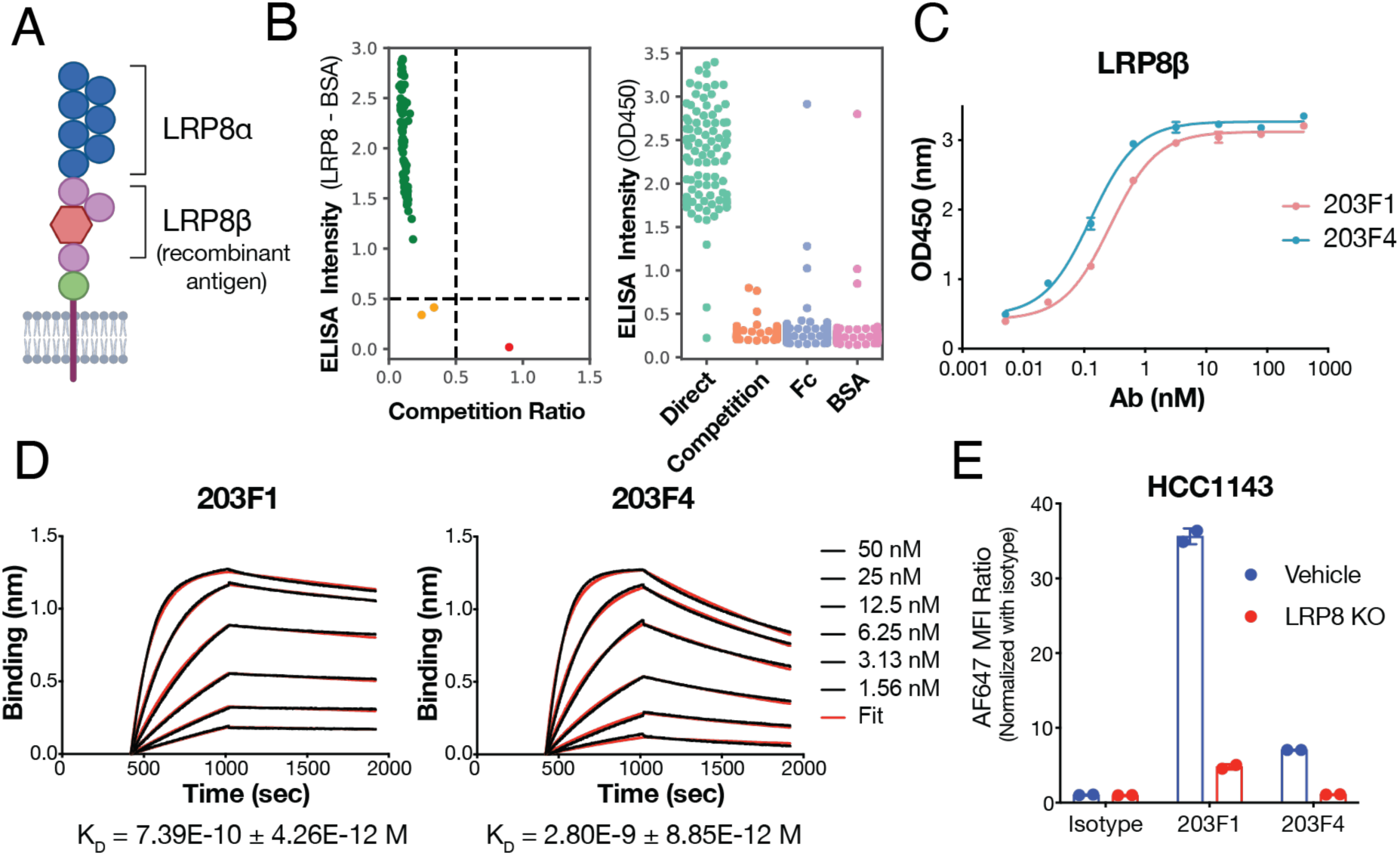
Generation and characterization of LRP8-specific antibodies. **(A)** The construct of full-length human LRP8, and the LRP8β was expressed as recombinant antigen with TEV-Fc-Avitag fusion for antibody selection by phage display. **(B)** Screening hits from phage selection. Plot on left shows screening phage for those that express well and are blocked from binding to antigen coated plate by soluble antigen in solution (upper left quadrant). Plot on right shows results from further testing in separate ELISA by direct antigen binding, competition with soluble antigen, direct binding to Fc, or BSA. **(C)** ELISA binding of purified recombinant selected Fabs against the LRP8 antigen. **(D)** BLI analysis of 203F1 and 203F4 Fabs against the recombinant LRP8β antigen. Biotinylated LRP8β was immobilized via the streptavidin biosensor and varying concentrations of each Fab were injected. Black lines are the experimental trace obtained from the BLI experiments and colored lines are the global fits. **(E)** Flow cytometry analysis of LRP8 IgG binding on HCC1143 control cells or LRP8 KO cells. Cells were stained with LRP8 antibodies for 30 min, followed by binding AF647 conjugated anti-human IgG. (H+L) antibody staining for 15 min. The ratio of mean fluorescent intensity (MFI) of the APC fluorescence channel was normalized to isotype control antibody. Data represents mean ± SD from two biological replicates.

After four rounds of phage selection, we plated 96 single phage colonies for screening using an enzyme-linked immunosorbent assay (ELISA)(32). Clones that specifically bound recombinant LRP8β-Fc but not Fc, BSA, or soluble LRP8β in competition assays (**Fig. 1B**) were sequenced and reformatted as recombinant fragment antigen-binding antibodies (Fabs). We identified four unique anti-LRP8 antibody clones, all of which bound MDA-MB-231 cells with endogenous LRP8 expression by flow cytometry and recombinant LRP8 by ELISA (**Figs. S1A-S1D**). None of the Fabs exhibited cross-reactivity with closely-related low-density lipoprotein receptor (LDLR), LRP2 or the N-terminal ligand binding domain of LRP8 (LRP8α) (**Figs. S1E-S1G**). All Fabs exhibited minimal polyreactivity against nonspecific antigen panels(33) (**Fig. S2A**), and were monodispersed by analytical size exclusion chromatography (**Fig. S2B**). Given that 203F1 and 203F4 clones displayed high affinities against LRP8β, we characterized them in further detail. ELISA assays also confirmed strong binding of both Fabs to LRP8β (**Fig. 1C**), with EC_50_ values of 0.19 nM and 0.28 nM, respectively. Biolayer interferometry (BLI) measurements demonstrated that 203F1 and 203F4 bound LRP8 with affinities of 0.74 and 2.8 nM, respectively (**Fig. 1D**). Epitope binning using BLI revealed that 203F1 and 203F4 could bind LRP8β simultaneously, indicating they recognize distinct, non-competing epitopes (**Fig. S3A**). Both antibodies selectively bound to LRP8-positive HCC1143 cells but not to LRP8 knockout (KO) cells, further supporting their specificity (**Fig. 1E, Figs. S3B-S3C**).

### LRP8 antibody blocks the uptake of selenoprotein P

To assess whether LRP8 antibodies interfere with SEPP1 binding, we expressed a recombinant cysteine-substituted variant of selenoprotein P (SEPP1-Cys), in which all ten selenocysteine (Sec) residues encoded by the UGA stop codon were replaced with cysteines to facilitate expression. BLI showed that SEPP1-Cys bound LRP8β with a high affinity of 0.61 nM (**Fig. 2A**). AlphaFold3 modeling of the SEPP1–LRP8 complex yielded a high-confidence interface (ipTM = 0.80), consistent with a stable interaction. Top-ranked models consistently predicted engagement of both SEPP1 domains with the LRP8 β-propeller, supported by Rosetta interface energy scoring (dG_separated = -53 REU) (**Fig. S4A**). Epitope binning revealed that 203F4 clone competed with SEPP1-Cys for binding whereas 203F1 did not (**Fig. 2B, Fig. S4B**).

**Fig. 2.**
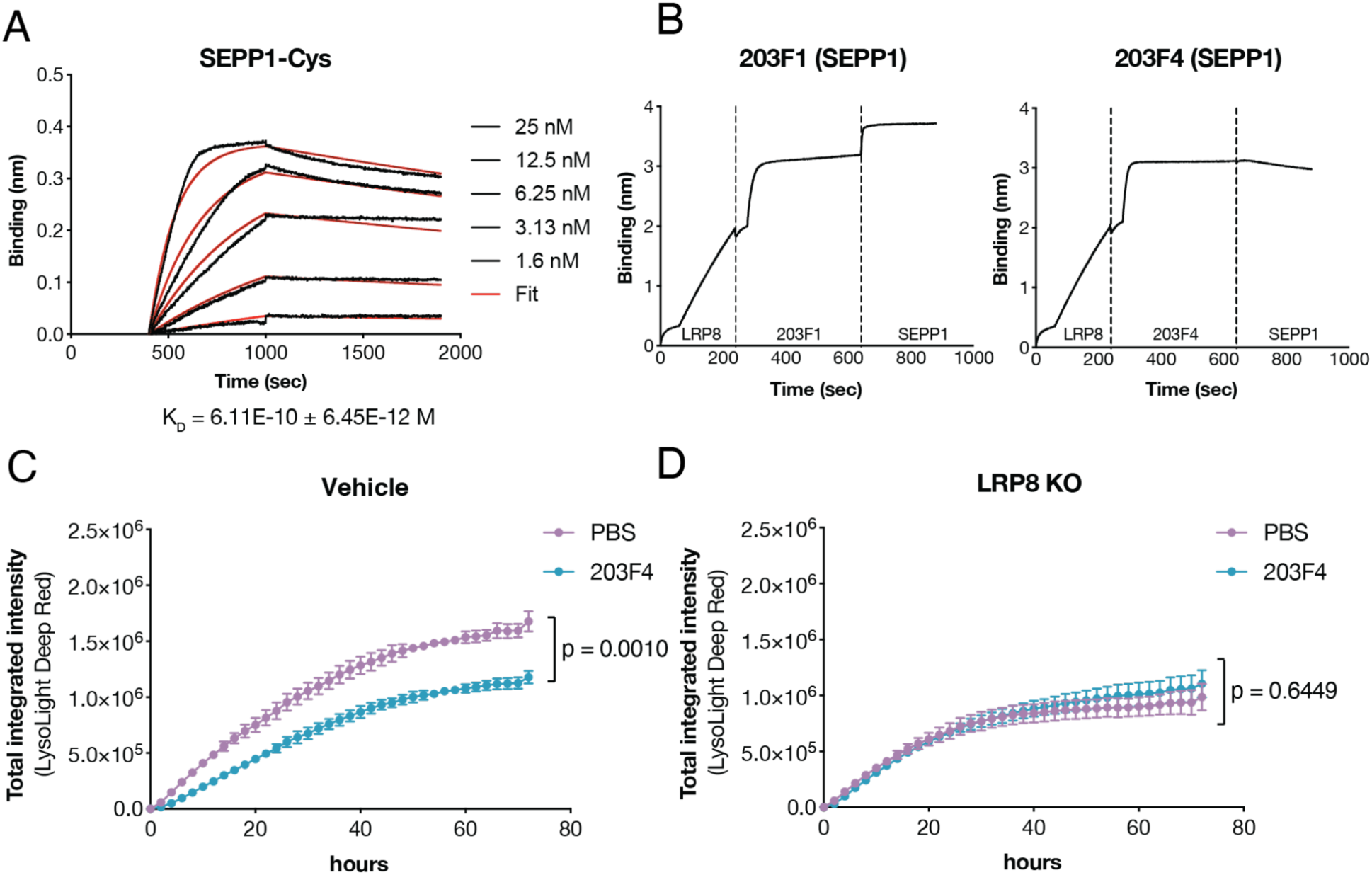
The 203F4 clone binds specifically to cells and blocks selenoprotein P binding. **(A)** BLI analysis of SEPP1-Cys against the recombinant LRP8β antigen. Biotinylated LRP8β was immobilized via the streptavidin biosensor and varying concentrations of SEPP1-Cys were injected. Black lines were the experimental trace obtained from the BLI experiments and colored lines were the global fits. **(B)** Epitope binning of SEPP1-Cys and LRP8 Fabs revealed two different epitopes on LRP8β. Biotinylated LRP8β was captured using a streptavidin biosensor and indicated Fabs at a concentration of 200 nM were incubated for 10 min followed by incubation with 50 nM of the competing SEPP1-Cys for 5 min. **(C-D)** Internalization of SEPP1-Cys in HCC1143 cells with control sgRNA (vehicle) or LRP8 sgRNA (knockout). SEPP1-Cys was labeled with the lysolight deep red dye (LLDR) to monitor lysosomal trafficking. Cells were pre-incubated with PBS or 200 nM 203F4 Fab for 30 minutes prior to 50 nM SEPP1-Cys-LLDR treatment. Images were captured every 2 h for 72 h on the Incucyte. Total integrated intensity was calculated by NIRCU x μm^2^/image. Data represents mean ± SD from four biological replicates. Statistics were calculated by one-way ANOVA and Holm-Sidak multiple comparison tests.

We then used an Incucyte-based assay to monitor the uptake of SEPP1. We conjugated SEPP1-Cys with the cathepsin-dependent fluorescent probe, LysoLight Deep Red (LLDR)(34), which generates fluorescent signals upon cleavage by lysosomal cathepsins. We serum starved the HCC1143 cells, pretreated with phosphate-buffered saline (PBS) or LRP8 antibodies, then added SEPP1-LLDR. Importantly, the 203F4 antibody significantly reduced SEPP1 uptake (**Fig. 2C**). The blocking activity is abolished with LRP8 KO cells, further indicating 203F4’s binding dependency on LRP8 (**Fig. 2D**). We reasoned that the residual internalization of SEPP1 may occur via macropinocytosis. To test this, we treated cells with Cytochalasin D(35), an actin polymerization inhibitor that suppresses membrane ruffling and macropinocytosis. Cytochalasin D almost completely abolished SEPP1 uptake, supporting the existence of a secondary macropinocytic pathway (**Fig. S4C**). In contrast, the 203F1 antibody had no effect on SEPP1 internalization, consistent with its recognition of a distinct, non-competitive epitope (**Fig. S4D**).

### KineTACs efficiently degrade LRP8

To selectively target LRP8 for degradation, we generated two bispecific LRP8-targeting KineTACs(27), with one CXCL12 arm recruiting the chemokine receptor CXCR7, and the other anti-LRP8 arm of 203F1 (termed KineTAC1) or 203F4 (termed KineTAC2), respectively (**Fig. 3A**). To avoid Fc effector functions, LALAPG mutations were introduced in both Fc chains(36).

**Fig. 3.**
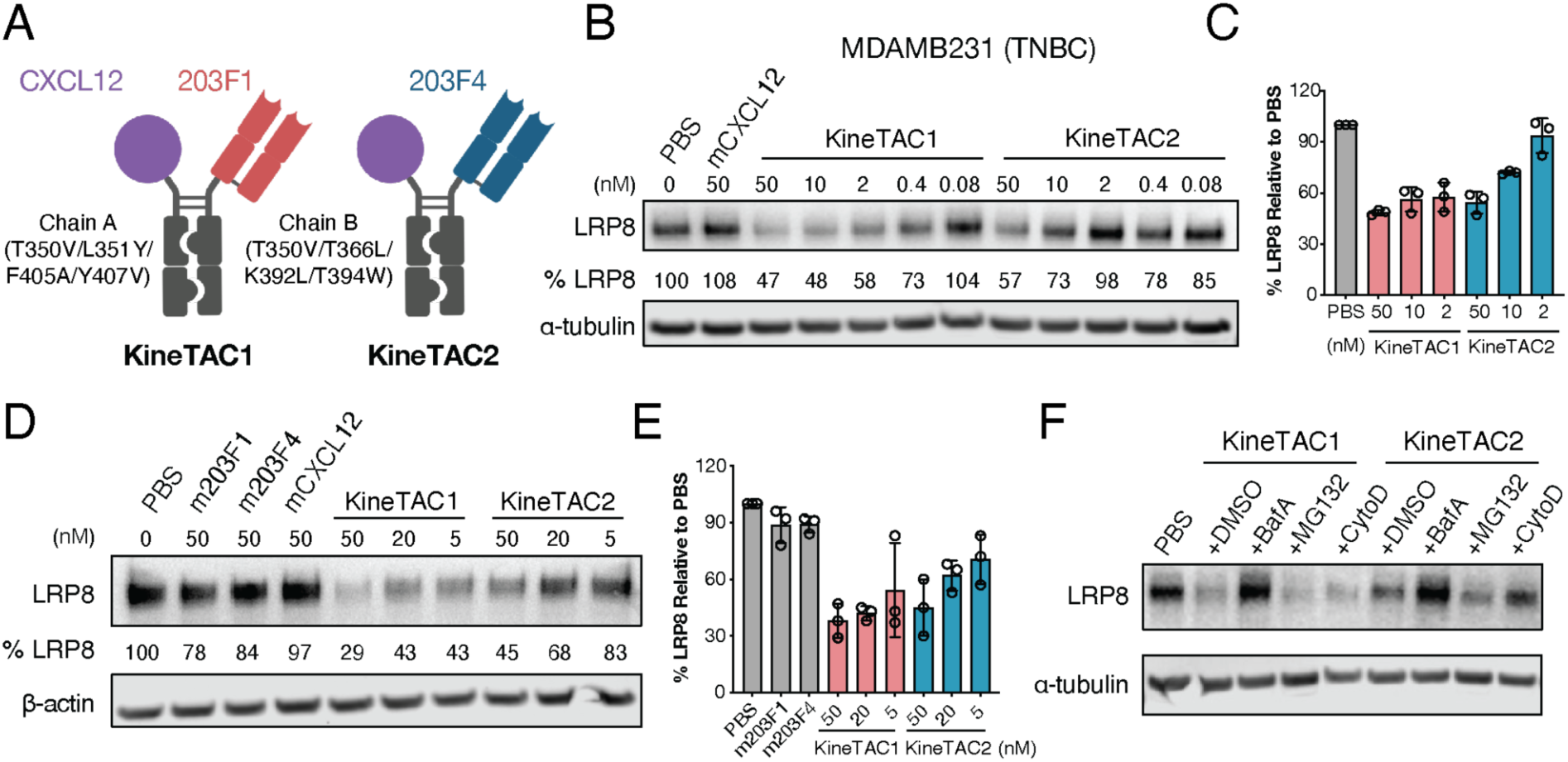
Generation of LRP8-degrading KineTACs. **(A)** Schematic illustration of bispecific LRP8-targeting KineTACs. Both KineTAC Fc chains contain LALAPG mutations to inactivate effector functions. **(B-C)** Western blotting showing degradation of LRP8 in triple negative breast cancer MDA-MB-231 cells following various concentrations of KineTAC treatment for 24 hours. Percent LRP8 levels were quantified by ImageJ relative to the PBS control. Data represents mean ± SD from three biological replicates. **(D-E)** Western blotting showing degradation of LRP8 in neuroblastoma SK-N-DZ cells following various concentrations of KineTAC or monomeric antibody treatment for 24 hours. Percent LRP8 levels were quantified by ImageJ relative to the PBS control. Data represents mean ± SD from three biological replicates. **(F)** Western blot analysis showing lysosome dependent LRP8 degradation on HCC1143 cells following KineTAC treatment. Cells were pretreated with either 500 nM Bafilomycin A (BafA), 500 nM MG132, or 500 nM Cytochalasin D (CytoD) for 1 h followed by 24 h treatment with 50 nM KineTAC1 or KineTAC2.

LRP8 is highly expressed in triple-negative breast cancer (TNBC) compared to other breast cancer subtypes and its genomic locus is amplified in 24% of TNBC tumors(37). In TNBC MDA-MB-231 cells, KineTAC1 induced robust LRP8 degradation within 24 hours, with activity detectable at concentrations as low as 2 nM (**Figs. 3B-3C**). By contrast, KineTAC2 was markedly less potent, requiring ∼50 nM for measurable degradation. The reduced activity likely reflects both weaker binding affinity of 203F4 and differences in epitope recognition. Similar results were obtained in TNBC HCC1143 cells and neuroblastoma SK-N-DZ cells, where KineTAC1 efficiently degraded LRP8 while KineTAC2 displayed reduced potency (**Figs. 3D-3E, Figs. S5A-S5B**). Monovalent LRP8 antibodies had no effect, indicating that degradation requires the CXCL12 arm for induced proximity. Given that LRP8 is an endocytic LDLR-family receptor, it may recycle to the plasma membrane upon antibody binding rather than undergoing self-degradation (38).

To further define the degradation mechanism, HCC1143 cells were pretreated with different pathway inhibitors prior to KineTAC exposure. LRP8 degradation was blocked by the lysosomal V-ATPase inhibitor, Bafilomycin A1(39), but not by the proteasome inhibitor MG132(40) or the macropinocytosis inhibitor Cytochalasin D (**Fig. 3F**, **Fig. S5C**). These findings confirm that KineTAC-mediated LRP8 degradation proceeds through a lysosome-dependent pathway, consistent with prior reports of extracellular protein degradation(27, 38).

### LRP8 degradation reduces selenoprotein levels

To determine whether LRP8 degradation affects selenoprotein translation, we treated HCC1143 cells with KineTAC1, KineTAC2, or the single armed monomeric antibodies for 48 hours. As expected, KineTAC1 efficiently degraded LRP8, whereas KineTAC2 was less potent and only effective at 50 nM (**Fig. 4A**). Both treatments reduced GPX4 abundance, although KineTAC2 was less effective, suggesting that active degradation of LRP8 has a stronger impact on selenoprotein translation than merely blocking ligand binding. Similar results were observed in HT-1080 fibrosarcoma cells, where KineTAC1 markedly reduced LRP8 and lowered GPX4 by ∼52% after 72 hours, while KineTAC2 again showed weaker effects (**Fig. 4B)**. Supplementation with sodium selenite or L-selenocystine restored GPX4 levels regardless of LRP8 degradation, indicating that reduced selenoprotein level arises from impaired selenium uptake (**Figs. S6A-S6B**).

**Fig. 4.**
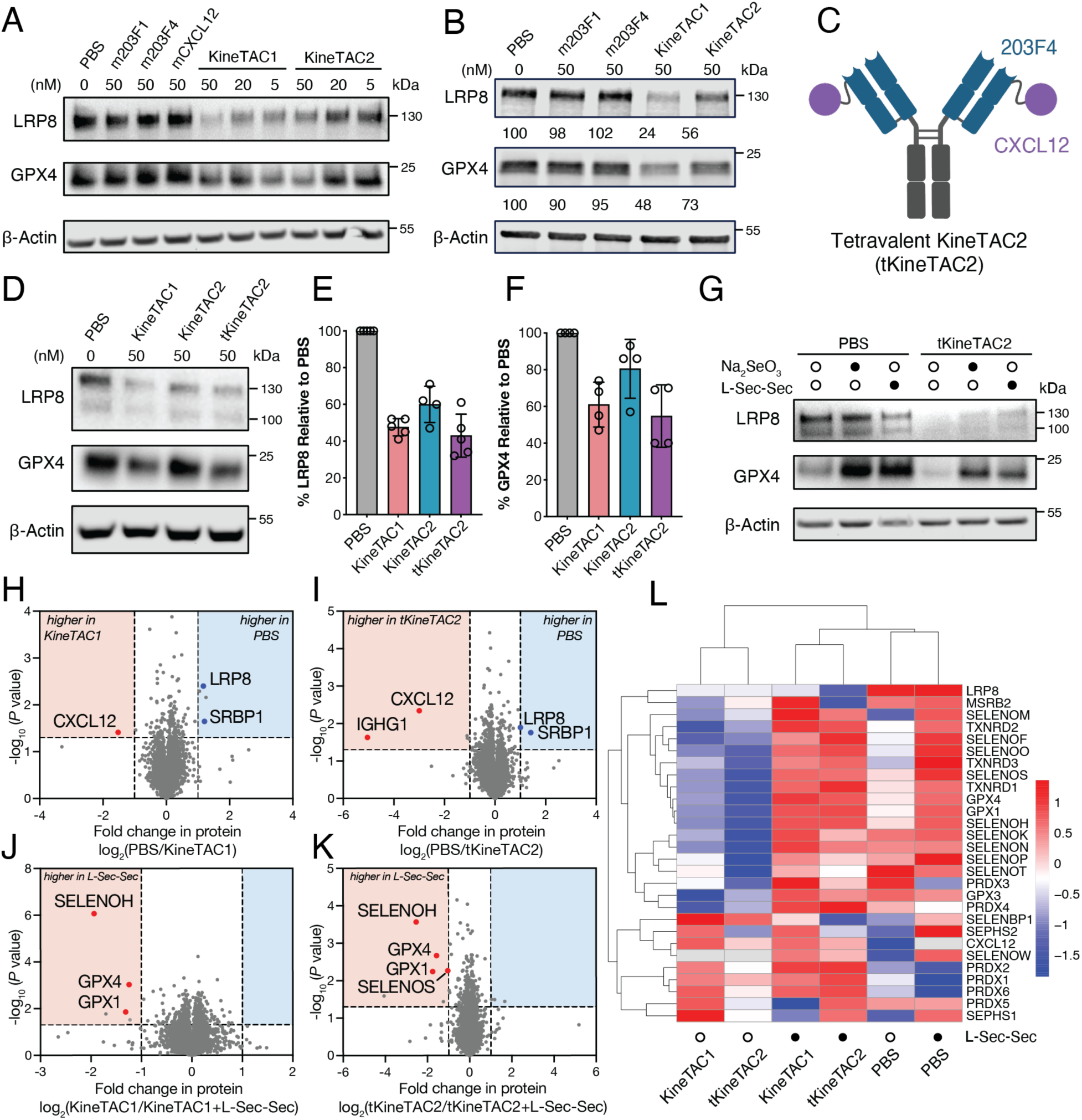
LRP8 degradation reduces GPX4 and selenoprotein translation. **(A)** Western blotting shows that KineTACs degrade LRP8 causing reduction in GPX4 levels in HCC1143 cells after 48 hours of KineTAC treatment, whereas treatment with monomeric antibodies have no such effect. Data is representative for at least three independent experiments. **(B)** Western blotting showing degradation of LRP8 and reduction of GPX4 in HT-1080 cells following 50 nM KineTACs treatment for 72 hours but not for monomeric antibodies (m203F1, m203F4). **(C)** Schematic of the tetravalent KineTAC2 (tKineTAC2). **(D-F)** Western blotting showing degradation of LRP8 and reduction of GPX4 in Kelly cells **(D)** following 50 nM KineTAC treatment for 72 hours. Quantification of LRP8 **(E)** and GPX4 **(F)** protein levels relative to a PBS control was performed using ImageJ. Data represents mean ± SD from four biological replicates. **(G)** Western blotting showing that selenium supplementation restores selenoprotein abundance following 72 hours of 50 nM tKineTAC2 treatment. GPX4 levels could be rescued by addition 50 nM sodium selenite and 50 nM L-selenocystine (L-Sec-Sec) despite ongoing LRP8 degradation. **(H-K)** Whole-cell proteomics of Kelly cells treated for 48 hours with PBS, 50 nM KineTAC1 **(H)**, 50 nM tKineTAC2 **(I)**, or the same treatments with 50 nM L-Sec-Sec supplementation **(J–K)**. Fold changes in protein levels were quantified by mass spectrometry. P values were calculated using two-tailed Student’s t-tests. **(L)** Heatmap showing abundance of selenocysteine-containing and related proteins in Kelly cells after 48 hours of treatment with PBS, KineTAC1, or tKineTAC2, with or without 50 nM L-Sec-Sec (filled dots and open dots, respectively).

Because the antibody 203F4 blocks SEPP1 uptake but lacks strong degradation activity, we sought to enhance avidity by fusing CXCL12 to the N-terminus of its light chain, generating a tetravalent KineTAC2 IgG (termed tKineTAC2) (**Fig. 4C, Fig. S6C**). This engineered antibody robustly degraded LRP8 and decreased GPX4 levels in Kelly cells, achieving greater degradation than KineTAC1 (**Figs. 4D-4F**), indicating that improved avidity helps to increase degradation activity(41). Consistently, inorganic selenium supplementation restored selenoprotein levels regardless of tKineTAC2-mediated LRP8 degradation (**Fig. 4G, Fig. S6D**).

To assess broader consequences of LRP8 degradation, we performed whole-cell proteomics in neuroblastoma Kelly cells after 48 hours of treatment with KineTAC1, tKineTAC2, or controls. LRP8 was the most significantly downregulated protein following KineTAC1 and tKineTAC2 treatment, underscoring the selectivity of KineTAC-mediated extracellular protein degradation(27) (**Figs. 4H, 4I, Fig. S7**). Interestingly, sterol regulatory element-binding protein 1 (SREBP1)(42, 43), a transcriptional factor that regulates lipid homeostasis, was also downregulated upon KineTAC treatment. It has been shown that LRP8 expression level is associated with lipid metabolism-related genes, including SREBP1(44). Consistent with western blot results, proteomics showed that L-selenocystine supplementation during KineTAC treatment rescued GPX4, GPX1 and other selenoproteins (**Figs. 4J, 4K**), confirming their dependence on extracellular selenium. Overall, we observed more downregulation of a panel of selenoproteins including the SELENO family, the thioredoxin reductase (TrxR) family, and the GPX family (**Fig. 4L**). Interestingly, the peroxiredoxin protein (PRDX) abundance, such as PRDX1 and PRDX2 did not change; these proteins contain cysteines instead of selenocysteines as antioxidant enzymes(45), further indicating the dependence of Sec in the LRP8-GPX4 axis (**Fig. 4L**).

Together, these findings show that LRP8 degradation selectively impairs selenium metabolism, resulting in reduced GPX4 and other selenoproteins, while also inducing broader metabolic alterations in cells.

### LRP8 degradation sensitizes tumor cells to ferroptosis

To determine whether disrupting the SEPP1–LRP8 axis enhances ferroptotic vulnerability, we treated HCC1143 cells with ferroptosis inducers in the presence of KineTACs. All KineTAC variants markedly sensitized cells to the covalent GPX4 inhibitor RSL3(13), and this effect was fully rescued by the radical-trapping antioxidant ferrostatin-1 (Fer-1)(46) (**Figs. 5A, 5B**). KineTAC1 and tKineTAC2 elicited stronger sensitization than KineTAC2, consistent with their ability to induce LRP8 degradation. KineTAC treatment also potentiated cell death induced by additional GPX4 inhibitors, ML162 and ML210 (**Figs. 5C, 5D**). To evaluate lipid peroxidation, HCC1143 cells were treated with 50 nM KineTACs for 72 h followed by RSL3 exposure. KineTAC1 and tKineTAC2 significantly elevated lipid peroxidation (**Figs. 5E, 5F**). Similar results were observed in Kelly cells, where all KineTAC treatments enhanced RSL3-induced cell death and lipid peroxidation (**Figs. 5G, 5H, Figs. S8A, S8B**). Notably, KineTAC treatment alone was sufficient to trigger lipid peroxidation in Kelly cells (**Fig. S8C**), highlighting the intrinsic ferroptotic hypersensitivity conferred by LRP8 loss in MYCN-amplified neuroblastoma(23).

**Fig. 5.**
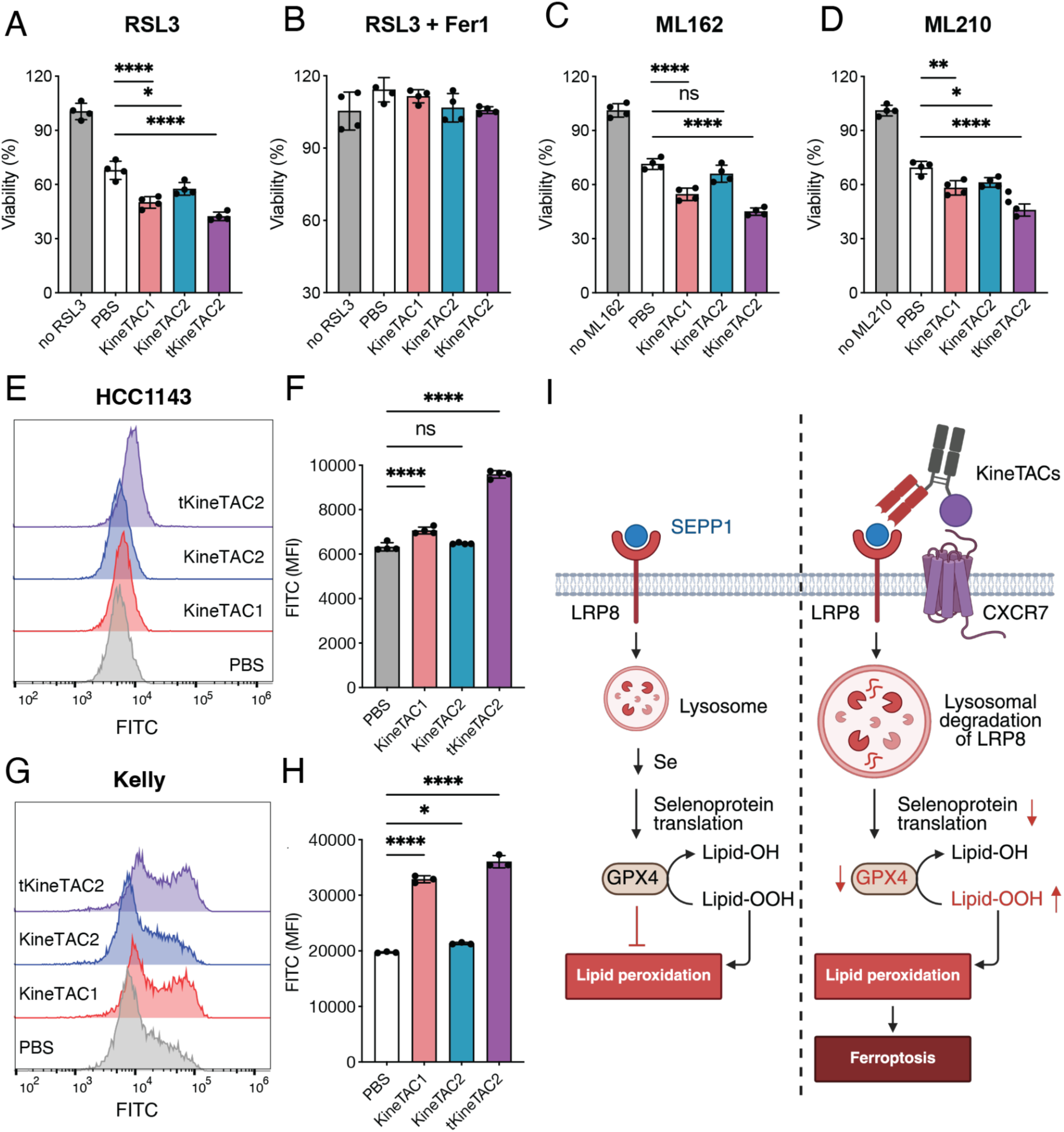
LRP8 degradation sensitizes tumor cells to ferroptosis. **(A-D)** Viability of HCC1143 cells treated with KineTACs for 48 h, followed by 24 h treatment with 1 μM RSL3 **(A)**, 1 μM RSL3 + 1 μM Fer-1 **(B)**, 200 nM ML162 **(C)**, or 1 μM ML210 **(D)**, respectively. Cell viability was measured using the CellTiter-Glo Reagent. Data represent quadruplicates ± SD. **(E-F)** Lipid peroxidation in HCC1143 cells pretreated with KineTACs for 72 h and then with 200 nM RSL3 for 5 h at 37 °C, detected using 5 μM BODIPY 581/591 C11 dye by flow cytometry. Gating was performed on singlets/live cells **(E)**, and GFP mean fluorescence intensity (MFI) was quantified **(F)**. Data represent quadruplicates ± SD. **(G-H)** Flow cytometry analysis of lipid peroxidation in Kelly cells pretreated with KineTACs for 72 h and then with 100 nM RSL3 for 5 h, detected using 5 μM BODIPY 581/591 C11 for 10 min at 37 °C, and counterstained with Near IR live/dead dye. Gating was performed on singlets/live cells **(G)**, and GFP MFI was quantified **(H)**. Data represent triplicates ± SD. **(I)** Schematic showing that CXCR7-mediated lysosomal degradation of LRP8 by KineTAC disrupts the LRP8–SEPP1 axis, reducing selenoprotein translation, elevating lipid peroxidation, and sensitizing tumor cells to ferroptosis. Schematic was generated using Biorender. All statistics were determined by one-way ANOVA. \**P* < 0.05. \*\**P* < 0.01. \*\*\*\**P* < 0.0001. ns, not significant.

We further used an Incucyte-based live-cell imaging assay to monitor ferroptotic susceptibility. Pretreatment with 50 nM KineTACs for 72 h sensitized Kelly cells to the system Xc⁻ inhibitor imidazole ketone erastin (IKE) (**Fig. S8D**), which indirectly compromises GPX4 function through depletion of cystine and glutathione. KineTAC treatment also enhanced sensitivity to RSL3 (**Fig. S8E**), consistent with the lipid peroxidation results. All ferroptosis phenotypes were fully rescued by Fer-1 (**Figs. S8D, S8E**). The differential magnitude of sensitization across ferroptosis inducers may reflect their distinct mechanisms of action: whereas IKE limits cystine uptake and depletes glutathione over time, RSL3 directly inactivates GPX4, leading to more acute but context-dependent lipid peroxidation responses. Differences in compound permeability, redox buffering capacity, and cell-type-specific antioxidant programs could also contribute to the observed variability.

Together, these results establish LRP8 degradation as a generalizable strategy to potentiate ferroptosis across tumor cell types. Mechanistically, targeted degradation of LRP8 disrupts the SEPP1–LRP8 axis, restricting selenium availability for selenoprotein biosynthesis, thereby perturbing lipid homeostasis and promoting lipid peroxidation (**Fig. 5I**).

## Discussion

Ferroptosis sensitivity in cancer is tightly constrained by access to key metabolic inputs, including selenium required for GPX4 synthesis. This dependence creates an opportunity to modulate ferroptosis indirectly by targeting extracellular pathways that control nutrient uptake rather than intracellular enzymes themselves. eTPD is emerging as a powerful modality for antibody-based therapeutics. In this study, we engineered antibodies recognizing two distinct epitopes on LRP8 and assembled them into bispecific LRP8-targeting KineTACs capable of robustly degrading the membrane bound LRP8. By selectively removing this receptor, we demonstrated that extracellular degradation can indirectly downregulate intracellular pathways that depend on selenium availability, providing direct evidence for the functional coupling of the LRP8–GPX4 axis. Whole-cell proteomics revealed the high selectivity of KineTAC-mediated degradation. Selenium supplementation restored GPX4 levels, further validating the selenium-dependent nature of this regulatory network. We also found that increasing molecular avidity and valency improves degradation potency, consistent with prior work on other degrading receptors such as folate receptors(41) and transmembrane E3 ligases(47).

Importantly, we show that targeting LRP8 sensitizes tumor cells to ferroptosis, highlighting LRP8 as a promising therapeutic entry point for regulating lipid peroxidation. In contrast, although GPX4 is a central suppressor of ferroptosis, it is also indispensable for normal tissue homeostasis. Systemic inhibition of GPX4 can therefore induce ferroptosis in healthy cells, including those in the kidney(14, 48), potentially narrowing the therapeutic window(49). Moreover, many early GPX4 inhibitors (e.g., RSL3, ML162, and ML210) are covalent compounds that perform well in cultured cells but display poor pharmacokinetics and limited target engagement *in vivo*(*50*). High chemical reactivity can promote promiscuous binding to proteins containing reactive cysteine or selenocysteine residues, leading to off-target cytotoxicity that is not directly attributable to ferroptosis.

Our findings suggest that extracellular targeting of LRP8 may offer a safer and more selective strategy to modulate ferroptosis, circumventing the liabilities associated with direct GPX4 inhibition. Certain cancers, particularly those with low system Xc⁻ activity, that contributes to the transport of specific selenium species, appear especially dependent on LRP8 and therefore more vulnerable than normal tissues. A prominent example is for MYCN-amplified neuroblastoma(23), in which tumor cells have limited alternative pathways for selenium uptake. Notably, our LRP8-directed KineTACs function both as receptor degraders and as functional SEPP1 blockers, thereby simultaneously reducing selenium uptake and compromising GPX4 expression/activity through a dual mechanism.

As an endocytic member in the LDLR family(51), LRP8 serves as a viral entry receptor (52–54) and potentially undergoes fast internalization, opening opportunities beyond protein degradation. The LRP8 antibodies we developed could be adapted into antibody-drug conjugates carrying GPX4 inhibitors(2, 55), enabling the selective delivery of ferroptosis-inducing agents into LRP8-expressing tumor cells. Together, these findings position LRP8 as a versatile therapeutic node and highlight KineTAC-based extracellular degradation as a compelling approach for sensitizing tumors to ferroptosis in a targeted and potentially safer manner.

## Acknowledgement

We are grateful to generous support from NIH-R01CA248323 (J.A.W), NIH-R35GM122451 (J.A.W.), the Hind Professorship in Pharmaceutical Sciences (J.A.W), and NIH-NCI R01CA276207 (J.A.O.). F.Z. is supported by an A.P. Giannini Foundation Fellowship. T.M.P.C. is supported by the National Cancer Institute of the National Institutes of Health (F32CA298768). A.I. is supported by The Pew Charitable Trusts and Chan Zuckerberg Initiative Foundation. M.K.C.S. is supported by the NSF Graduate Research Fellowship Program (2445150).

## Author Contributions

F.Z. and J.A.W. conceived and designed the study. F.Z., M.K.C.S. performed phage selection, F.Z. performed antibody screening, internalization and characterization experiments. F.Z., S.G., Y.Z., and Y.W., cloned and expressed the recombinant proteins. F.Z., A.I. performed the western blotting experiments. F.Z., T.M.P.C., and Y.C. performed the proteomics and processed the data. A.I and J.A.O. provided the LRP8 KO cells. F.Z. and A.I. performed ferroptosis cell death assays. K.K.L provided the Rosetta energy analysis. F.Z., J.A.O., and J.A.W. wrote the manuscript and all authors reviewed and edited the manuscript.

## Materials and Methods

### Plasmid construction

All the IgGs were constructed in a pcDNA3.4 vector that expresses the light chain and heavy chain, respectively for mammalian expression. For generating bispecific antibodies, the heavy chain variable regions and CXCL12 were cloned into zymework-A mutant Fc (T350V/L351Y/F405A/Y407V) and zymework-B mutant Fc (T350V/T366L/K392L/T394W) sequences respectively(56). Both Fc chains contained additional L234A/L235A/P329G mutations(36) to eliminate Fc-effector function for macrophage and NK cell recruitment. For generating the tetravalent KineTAC2, CXCL12 was cloned into the N-terminus of 203F4 light chain with (G_4_S)_3_ linkers whereas 203F4 heavy chain was cloned in unmutated Fc. LRP8β antigen was cloned into pFuse vector with IL-2 signal peptide followed by tobacco etch virus (TEV) protease, Fc, and Avitag sequence in the C-terminus. The 10 selenocysteines from SEPP1 (Uniprot P49908) were replaced with cysteines, and then cloned into the pcDNA3.4 vector with a C-terminal His10 tag. All the Fabs were constructed in a dual-expression pBL347 vector that expresses the light chain and the heavy chain with the pelB and the stII signal peptides, respectively, for the periplasm expression.

### Cell lines

MDA-MB-231 cells were cultured in DMEM (ThermoFisher Scientific) with 10% fetal bovine serum (FBS) and 1% penicillin–streptomycin. HCC1143, Kelly, and HT-1080 cells were cultured in RPMI-1640 (ThermoFisher Scientific) with 10% FBS and 1% penicillin–streptomycin. HCC1143 LRP8 vehicle (Cas9) cells and KO3(24) cells were supplemented with 100 μg/mL hygromycin (Sigma-Aldrich). Expi293 cells were cultured in Expi293 expression medium (ThermoFisher Scientific).

### Phage display

Phage selection was done as described previously(32). In brief, library E from University of Chicago and UCSF library were incubated with streptavidin-coated magnetic beads pre-conjugated with biotinylated Fc protein to remove nonspecific binders. Unbounded phage were then incubated with streptavidin-coated magnetic beads pre-conjugated with biotinylated LRP8β-TEV-Fc antigens. After 4 washes, antigen-bound phage were eluted from beads by incubating with 1 μM TEV protease for 20 min. In total, four rounds of selections were performed with a decreasing concentration of LRP8β-TEV-Fc-antigen (1000, 50, 20, 10 nM). From round 3, the phage library was first enriched by protein A magnetic beads to deplete nondisplayed or truncated Fab phage before each round of the selection.

### Phage ELISA

384-well Maxisorp plates were coated with Neutravidin (10 μg/mL) overnight at 4 °C and subsequently blocked with BSA (2% w/v) for 1 h at RT. 20 nM biotinylated LRP8β-TEV-Fc, BSA, or empty Fc antigens were captured on the NeutrAvidin-coated wells for 30 min followed by the addition of 1:5 diluted single-colony phage for 1 h. For competition, 20 nM of soluble LRP8β-TEV-Fc antigen was added to the diluted phage and incubated for 1 h. The bound phage was then detected by a horseradish peroxidase (HRP)-conjugated anti-M13 phage antibody (Sino Biological). The ELISA plates were washed three times after each incubation, and antibody binding was detected by TMB substrate (VWR) and read at 450 nm.

### Protein expression

IgGs and antigen were expressed in Expi293 cells. For 30 mL transfection, 24 μg of DNA was added to 3 mL of OptiMEM, followed by 24 μL of FectoPro transfection reagents. After 10 min of incubation, 27 mL of Expi293F cells at 3 millions/mL were added and shake at 37°C. Biotinylated antigens were expressed in Expi293 cells stably expressing BirA, and 3 mL of 4 mM biotin was added at transfection. On the second day, 300 μL of 300 mM vaporic acid and 270 μL of 45% glucose were added to the cells. After 5 days, cultures were harvested, centrifuged at 4,000 g for 20 min, and clarified through a 0.45-μm Steriflip filter. Supernatants were then incubated with Sepharose A resin (Cytiva) for 2 h, proteins were then eluted by 0.1 M acetic acid and neutralized by Tris pH 11. Proteins were buffer-exchanged at least twice into PBS using Amicon concentrators (Millipore Sigma). For His-tagged proteins, culture supernatants were incubated with HisPur™ Ni-NTA resin (Thermo Fisher Scientific) for 2 h at 4 °C and eluted with 500 mM imidazole, followed by PBS buffer exchange using Amicon concentrators. Fabs were expressed in *Escherichia coli* C43 (DE3) Pro+ grown in an optimized TB autoinduction medium at 37 °C for 6 h, cooled to 30 °C for 18 h. Cells were harvested by centrifugation and lysed using B-PER lysis buffer (ThermoFisher Scientific). The lysate was incubated at 60 °C for 20 min and centrifuged to remove the inclusion body. The Fabs were purified by Sepharose A resin via affinity chromatography and buffer exchanged in PBS for further characterization. Purity and integrity of all proteins were assessed by SDS–PAGE (Thermo Fisher Scientific).

### Recombinant protein ELISA

384-well Maxisorp plates were coated with neutravidin (10 μg/mL) or 6x Histag antibody (Invitrogen, Cat# MA1-21315, 2 μg/mL) overnight at 4 °C and subsequently blocked with BSA (2% w/v) for 1 h at RT. 20 nM of antigens were captured onto pre-coated wells for 1h. Recombinant LDLR, LRP2, and LRP8 proteins were purchased from ACROBiosystems. For polyspecificity ELISAs, the autoantigens cardiolipin (Sigma, 50 μg/mL), insulin (Sigma, 1 μg/mL), lipopolysaccharide (LPS, InvivoGen, 10 μg/mL), and single-stranded DNA (ssDNA, Sigma, 1 μg/mL), were directly coated onto plates overnight 4 °C. After three washes, serially diluted Fabs were added to the plates and incubated for 1 h at RT. After three washes, 1:5000 diluted peroxidase-anti-human IgG (H+L) (Jackson ImmunoResearch) were added to the plates and incubated for 30 min. After three times of washing, antibody binding was detected by TMB substrate (VWR), quenched by 1 M phosphoric acid, and read at 450 nm.

### Biolayer interferometry

BLI experiments were performed at room temperature using an Octet RED384 instrument (ForteBio). 20 nM biotinylated antigen-Fc was immobilized to an optically transparent SA biosensor (ForteBio). Different concentrations of monomeric Fab antibodies or SEPP1-Cys in kinetics buffer (PBS, 0.05% Tween-20, 0.2% BSA) were used as the analyte in a 384-well microplate (Greiner Bio-One). Because the Fc format presents antigens bivalently on the sensor surface, affinities (K_D_s) were best fit with a 1:1 binding model and calculated by a global fit analysis using the Octet RED384 Data Analysis HT software.

### Epitope binning by BLI

Anti-LRP8 antibodies were binned into epitope specificities using an Octet RED384 system. 20 nM of biotinylated LRP8β-Fc antigens were captured using streptavidin biosensors (Fortebio). After antigen loading, a saturating concentration of antibodies or SEPP1 (200 nM) was added for 10 min. Competing concentrations of antibodies or SEPP1 (50 nM) were then added for 5 min to measure binding in the presence of saturating antibodies. All incubation steps were performed in PBS/0.05% Tween-20/0.2% BSA.

### SEPP1 uptake assay

SEPP1-Cys was labeled using the LysoLight Antibody Labeling Kits (Invitrogen, Cat# L36003) following the manufacturer’s instructions. Briefly, SEPP1 were labeled with LLDR with a molar ratio of 1:6 in the presence of 100 mM sodium bicarbonate (pH 8.4) for 2 h at RT. Labeled SEPP1 was then purified with 7k Zeba dye and biotin removal columns (Thermo Scientific, Cat#A44297). HCC1143 cells were seeded at 5000 cells/well on a 96-well polystyrene tissue culture treated plate (Corning, Cat#3596). The next day, the media was removed and cells were serum starved for 12 h. Next, cells were pretreated with 200 nM anti-LRP8 Fabs or PBS for 30 min at 37 °C respectively, followed by 50 nM of SEPP1-LLDR treatment with or without 0.5 μM cytochalasin D. Cells were then imaged by Incucyte (Sartorius) every 2 h. Internalization was calculated by total integrated intensity (NIRCU x μm^2^/image) on the Incucyte software.

### Structural modeling and energy analysis

Structural models of the SEPP1–LRP8 complex were generated using AlphaFold3 (https://alphafoldserver.com/). The amino acid sequences of human LRP8β and SEPP1 were used as input, with selenocysteine residues in SEPP1 substituted with cysteine to enable modeling. Default multimer settings were applied, and top-ranked models were selected based on predicted alignment error (PAE) and confidence scores. Predicted complexes were visualized and analyzed using ChimeraX. To evaluate binding energetics, AlphaFold-derived models were subjected to Rosetta-based scoring (Rosetta 3.71). Structures were relaxed using the FastRelax protocol, and interface energies (ΔG) were calculated using the InterfaceAnalyzer module with default parameters. Residue-level contributions and interface metrics were extracted to assess key determinants of SEPP1–LRP8 binding.

### Flow cytometry

Cells were harvested by 0.05% Trypsin-EDTA and centrifuged at 400 g for 5 min. Pellets were washed once with PBS + 1% BSA. Cells were incubated with Fab or IgG antibodies in PBS + 1% BSA for 30 min at 4 °C. After two washes, cells were then stained with Alexa fluor 647 goat anti-human IgG (H+L) (Invitrogen, Cat#A-21445, 1:1000) for 30 min at 4 °C. After two washes, cells were resuspended in PBS + 1% BSA. Flow cytometry was performed using a CytoFLEX cytometer (Beckman Coulter, v.2.3.1.22) and CytoExpert software (v.2.3.1.22). Data were analyzed with FlowJo (v.10.8.0).

### Chemicals and reagents

1S,3R-RSL3 (referred to as RSL3) (Cat#HY-100218A) and MG132 (Cat#HY-13259) were purchased from MedChemExpress. Cytochalasin D (Cat#11330), ML162 (Cat#20455), ML210 (Cat#23282), IKE (Cat#27088), Ferrostatin-1 (no. 17729), FINO2 (Cat#25096), were all purchased from Cayman Chemical Company. Bafilomycin A1 (Cat#sc-201550A) was purchased from Santa Cruz Biotechnology. L-Selenocystine (Cat#545996), sodium selenite (Cat#S5261), acetonitrile (HPLC grade), H2O (HPLC grade), and formic acid (LC-MS grade) were obtained from Sigma-Aldrich.

### Degradation experiments

Cells were plated onto 6- or 12-well plates (Corning) and grown to ∼70% confluency before treatment. On the next day, cell culture medium was aspirated, various concentrations of antibodies in 1-2 mL of culture medium were then added to each well. Cells were incubated for 24 h or 48 h or 72 h at 37 °C prior to western blotting experiments. For inhibitor assays, cells were pretreated with 0.5 µM Bafilomycin A, 0.5 µM MG132, or 0.5 µM Cytochalasin D for 1 h, respectively prior to antibody treatment.

### Western blotting

Cells were lifted with versene (ThermoFisher Scientific), washed twice with PBS, and then lysed in RIPA buffer (EMD Millipore) with cOmplete mini protease inhibitor cocktail (Sigma-Aldrich) for 20 min on ice. The cell lysates were then centrifuged at 20,000 g for 10 min at 4 °C to remove any cell debris. Soluble protein concentrations were quantified by Rapid Gold BCA Protein Assay Kit (Pierce). Lysates were mixed with 4× Nupage LDS Sample Buffer (Invitrogen) and 2-mercaptoethanol, and then run on NuPAGE™ 4-12% Bis Tris Protein Gels (ThermoFisher Scientific). Proteins were transferred to polyvinylidene difluoride membranes using the iBlot2 Western Blotting Transfer System (Thermo Scientific). Membranes were blocked with TBS + 5% BSA + 0.5% Tween (PBST) for 30 min, and stained with primary antibodies overnight. After three washes, membranes were stained with secondary antibodies for 1 h at RT. After three washes, membranes were imaged with a LICOR imager (LI-COR Biosciences) or the ChemiDoc MP imaging system (BioRad). For HRP detection, Supersignal West Pico PLUS Chemiluminescent Substrate (Thermo Scientific, Cat# 34577) was added to the membrane. Antibodies used included rabbit anti-human ApoER2 (Abcam, Cat#ab108208, 1:1,000), rabbit anti-human GPX4 (Abcam, Cat#ab125066, 1:2,000), rabbit anti-human GPX1 (Abcam, Cat#ab22604, 1:1000), mouse anti-human SELENOM (Santa Cruz, Cat#sc-514952, 1:1000), mouse anti-human TrxR1 (Santa Cruz, Cat#sc-28321, 1:1000), mouse anti-human α-tubulin (Cell Signaling Technology, 3873S, 1:3,000), mouse anti-human β-actin (Cell Signaling Technology, 3700S, 1:3,000), IRDye 800CW goat anti-rabbit IgG (LI-COR Biosciences, Cat#926-32211), IRDye 680RD goat anti-mouse IgG (LI-COR Biosciences, Cat#926-68070, 1:5000), peroxidase goat anti-rabbit IgG (H+L) (Jackson ImmunoResearch, Cat# 111-035-144, 1:5000).

### Proteomics sample preparation

0.5 M Kelly cells were seeded onto 6-well plates in RPMI1640 medium supplemented with 10% dialyzed FBS and 1% penicillin–streptomycin overnight at 37 °C. The next day, media were removed, and 50 nM of antibodies were added to cells with or without 50 nM of L-selenocystine. After 48 h of treatment, cells were lifted with versene, and transferred to protein LoBind Eppendorf tubes. Cell pellets were washed three times with cold PBS, and flashed frozen until proteomic sample preparation.

Proteins were digested and desalted using the Preomics iST kit according to the manufacturer’s specifications (PreOmics, Cat#P.O.00027). Cell pellets were resuspended in the LYSE solution and sonicated with probe at 20% amplitude for 1 min (2 s on/4 s off). Cell lysates were clarified at 16,000 x g for 10 min and supernatant was incubated at 95 °C for 10 min for cysteine reduction and alkylation. DIGEST enzyme pellets were reconstituted in 210 µL RESUSPEND solution. Samples were cooled to room temperature and combined with 50 µL DIGEST solution. Incubation was performed at 37 °C for 4 hr while shaking. After the digestion, quenching of enzymatic activity was performed by adding 100 µL STOP solution to digested samples to a final pH below 2.0. Samples were spun through a PreOmics desalting column at 3,800 x g for 2 min. Samples were then washed with 200 µL WASH 1 and 200 µL WASH 2 each at 3,800 x g for 1 min. Peptides were eluted using 100 µL ELUTE solution in a new low-bind tube at 3,800 x g for 2 min. Desalted samples were dried down via SpeedVac and resuspended in 0.2% formic acid (FA) in water. Peptide concentration was determined using a Pierce Quantitative Colorimetric Peptide Assay (Thermo Fisher Scientific, Rockford, IL) before liquid chromatography–tandem mass spectrometry analysis.

### Liquid chromatography

Liquid chromatography was performed using either a Vanquish Neo UHPLC system (Thermo Fisher Scientific), configured in a direct injection format, or using a nanoElute 2 UHPLC (Bruker), configured in a trap-and-elute format. Peptides were separated on the Vanquish Neo with an Aurora Ultimate C18 120 Å, 1.7 µm, 75 µm x 25 cm UHPLC column (Ion Opticks). During LC separations, mobile phase A (MPA) was 0.1% FA in water and MPB was 80% ACN in water with 0.1% FA. For profiling of the proteome, a 90-min gradient ramped, at a flow rate of 250 nL/min, from 0-1% MPB from 0 – 1 min, 1-28% MPB from 1 – 80 min, 28 – 58% MPB from 80 – 89 min, 58 – 99% MPB from 89 – 90 min, and held at 99% MPB to 100 min before the column was washed and re-equilibrated at 0% MPB for three column volumes. During peptides separations, the LC column was held at 50 °C. On the timsTOF platform, peptides were separated using a nanoElute 2 with a 100 Å, 5 μM, 5 mm x 0.3 mm trap cartridge (Thermo Fisher) in line with a PepSep C18 100 Å, 1.5 μM, 25cm x 150 μM column on a timsTOF Pro 2 with a Captive Spray source and a nanoElute line (Bruker, Hamburg, Germany). During LC separations, MPA was 0.1% FA in water and MPB was ACN with 0.1% FA. A 62-min gradient ramped, at a flow rate of 500 nL/min, from 2-28% MPB from 0 – 60 min, 28-32% MPB from 60 – 62 min, 32 – 95% MPB from 62 – 62.5 min, and held at 95% MPB to 70 min before the column was washed and re-equilibrated at 0% MPB for 8 min. During peptide separations, the LC column was held at 50 °C.

### Mass spectrometry

Data independent acquisition (DIA) was performed either on a Thermo Scientific^TM^ Orbitrap Eclipse^TM^ Tribrid^TM^ hybrid mass spectrometer system (Thermo Fisher Scientific, San Jose, USA) or on a timsTOF Pro 2 (Bruker, Hamburg, Germany). On the Orbitrap Eclipse, precursors were ionized using electrospray ionization at 2 kV with respect to ground. The inlet capillary was held at 275 °C, and the ion funnel RF was held at 60%. During DIA experiments, all MS1 survey scans were acquired at a resolving power of 60,000 in the Orbitrap analyzer with a scan range of m/z 380 – 980, maximum injection time of 50 ms, and AGC target of 1,000,000 charges. DIA scans were acquired in the Orbitrap analyzer at resolution 15,000 with precursor mass range set to m/z 380-980. An isolation width of 14 Th with 1 Th window overlap created 42 bins across the m/z range. HCD normalized collision energy (NCE) was set to 30%. MS/MS scans required an injection time of 22 ms or 100,000 charges. For data acquisition on the timsTOF platform, ions were generated in positive mode with the capillary set to 4500 V. diaPASEF windows were generated between 0.69 V•s/cm2 to 1.29 V•s/cm2 in the 1/K0 dimension and between m/z 273 to 1173 in the mass dimension. A mass width of 25 m/z was used, yielding 36 steps per cycle. Ramp time was set to 120 ms, ramp rate was 7.93 Hz, and cycle time estimate was 1.26 s. A scan range of m/z 100 to 1700 was used.

### Data analysis

DIA .raw and .d files were processed using Spectronaut (20.2.250922.92449). Settings denote proteolytic cleavage C-terminal to lysine and arginine not before proline with up to 2 missed cleavages. A FASTA file containing the reviewed human proteome and isoforms was downloaded from UniProt on April 9, 2024 and imported into Spectronaut, where mutation-based decoy sequences were generated for false discovery rate estimation. Peptide length of 7 to 52 amino acids was specified. Alkylation of cysteines was fixed while oxidation of methionine and protein N-terminal acetylation were variable modifications. Data was processed in library-free directDIA mode. Precursors, peptide identifications, and proteins were filtered to maintain 1% FDR. Proteins identified with only one peptide were filtered out of the final dataset. Data were further analyzed in RStudio (2025.09.2) using Limma differential analysis.

### BODIPY 581/591 C11 assay

Cells seeded in a 12-well plate were treated with KineTACs or controls for 24 h 37°C, followed by 200 nM RSL3 treatment for 5 h and washed once with DPBS containing calcium and magnesium. Cells were detached from the plate with versene and incubated with 5 μM BODIPY 581/591 C11 (Invitrogen, Cat#D3861) at 37°C for 10 min. After two washes, cells were then stained with LIVE/DEAD™ fixable NIR-IR dye (Invitrogen, Cat# L10119). After two washes, cells were resuspended in PBS + 1% BSA. Flow cytometry was performed using a CytoFLEX cytometer (Beckman Coulter, v.2.3.1.22) and CytoExpert software (v.2.3.1.22). Data were analyzed with FlowJo (v.10.8.0).

### Cell viability and death analysis

For cell viability analysis, the CellTiter-Glo assay (Promega) was performed according to the manufacturer’s instructions. Briefly, 5000 cells/well were seeded on a 96-well polylysine-coated white plate (Corning, Cat#3917). The next day, media was aspirated, 100 μL fresh medium containing 100 nM of KineTACs or control antibodies were added to the cells for 24 h at 37 °C. Then 100 μL of ferroptosis compounds at various concentrations were added to the cells, with or without 1 μM Fer1 (Cayman Chemical Company). After 24 h of compound treatment, cell culture media was removed and viability was measured using CellTiter-Glo Reagent (Promega). The plates were incubated at room temperature for 10 min, and the luminescent signal was measured on the Spark microplate reader (Tecan).

Cell death was also analyzed using the IncuCyte S3 live-cell imaging system (Essen Bioscience) following a modified protocol adapted from Zhang *et a*l(57). Prior to treatment with ferroptosis-inducing agents (RSL3 or ML-210), Kelly cells were seeded at 1,500 cells per well in triplicate in black 384-well plates (Greiner, Cat. #781091) and pretreated with 50 nM KineTAC1, 50 nM tKineTAC2, or PBS for 72 h in an incubator set to 37 °C and 5% CO₂.

For Kelly cells, a separate drug plate was prepared by titrating different concentrations of RSL3 or ML-210 in a low-adhesion 384-well plate (PlateOne, Cat. #1884-2410). Medium containing 60 nM SYTOX Green Dead Cell Stain, with or without 2 µM Fer-1, was added to the corresponding wells. Plates were homogenized using a Bioshake 3000 ELM orbital shaker at 2,400 rpm for 45 s. The drug-containing medium was then carefully transferred onto the pre-seeded cells (black 384-well plates), resulting in final concentrations of 30 nM SYTOX Green and 1 µM Fer-1. Plates were immediately placed in the IncuCyte S3 imaging system within an incubator maintained at 37 °C and 5% CO₂. Phase-contrast and green fluorescence images were acquired every 2 h over 24- or 72-h periods. Cell death was quantified as the ratio of SYTOX Green–positive objects per image to total cell area (confluence %) using IncuCyte S3 image analysis software (Essen Bioscience). For each treatment condition, these ratios were plotted over time, the area under the curve (AUC) was calculated, and mean AUC values were plotted as a function of drug concentration using Prism (GraphPad).

