## Supplemental File for "Targeted extracellular degradation of LRP8 promotes ferroptosis in cancer cells"

**A**

| Ab ID | LC3 | HC1 | HC2 | HC3 |
|---|---|---|---|---|
| 203F1 | FGWSYLI | LSYYSI | SIYPYSGYTS | DHRYYDYAYYPGRYWGGF |
| 203F2 | GYYLI | LSYSSM | SIYSSYGYTS | GWASWDYYAARYGM |
| 203F3 | YWSYAPF | VSYSSI | SIYPYSGSTY | HEWYGSYYYGYYQYGL |
| 203F4 | YSSYSHYLI | ISSSYI | SIYSYYGSTY | HGAM |

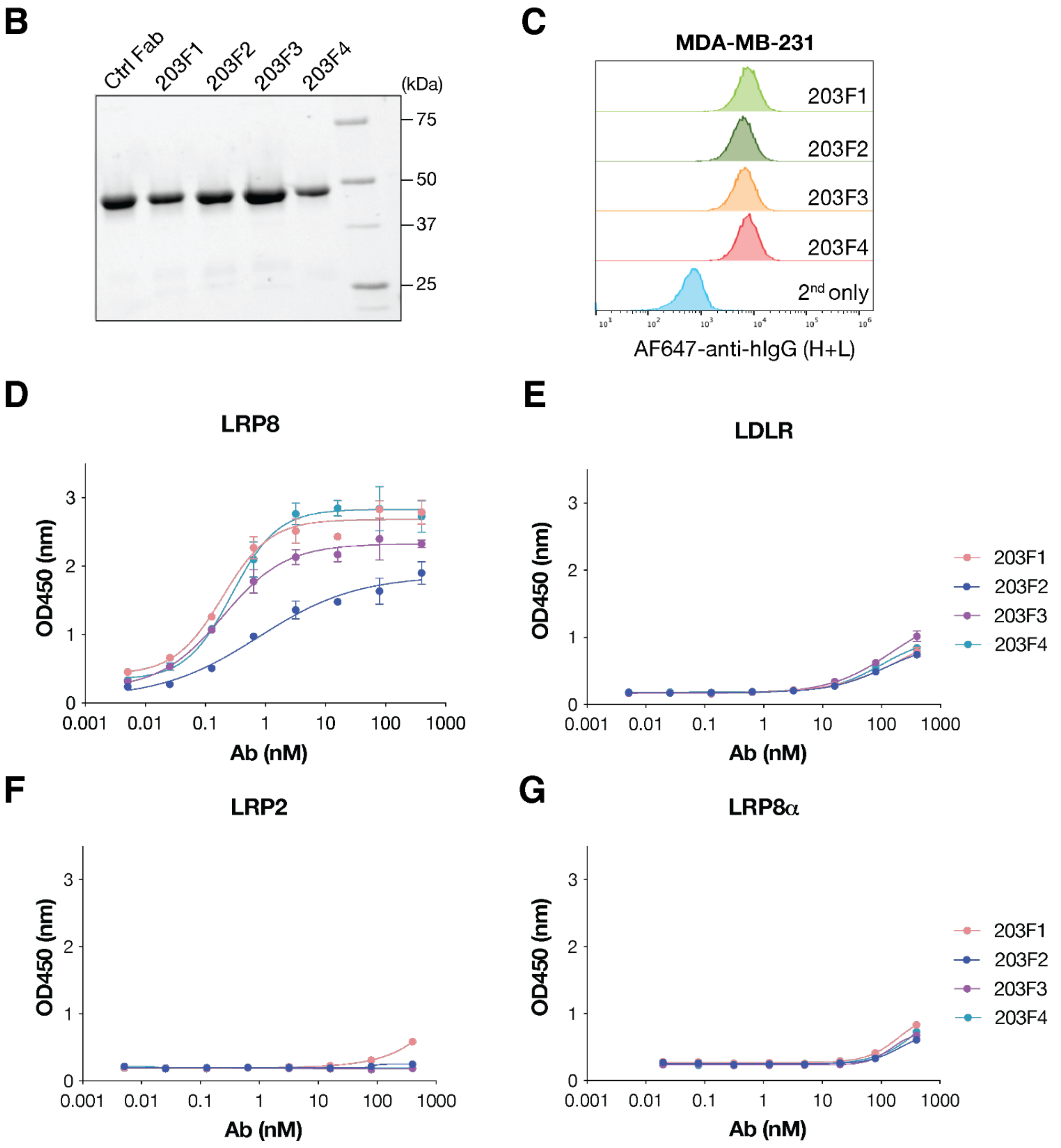

**Fig. S1**. **Characterization of LRP8-specific antibodies. (A)** Complementarity-determining region (CDR) sequences of LRP8 antibody heavy chain and light chain. **(B)** SDS-PAGE analysis of recombinant LRP8 Fabs. **(C)** Flow cytometry analysis revealed binding of Fabs against MDA-MB-231 cells with endogenous LRP8 expression. Cells were stained with 20 nM LRP8 Fabs for 30 min, followed by AF647 conjugated anti-human IgG (H+L) antibody staining for 15 min. Data is representative from two independent experiments. **(D-G)** ELISA binding of LPR8 Fabs against recombinant **(D)** LRP8, **(E)** LDLR, **(F)** LRP2, and **(G)** LRP8α. Binding was detected by peroxidase-conjugated goat anti-human IgG (H+L) antibodies. Each sample was tested in duplicate and repeated for at least two independent experiments. Error bars represent standard deviation.

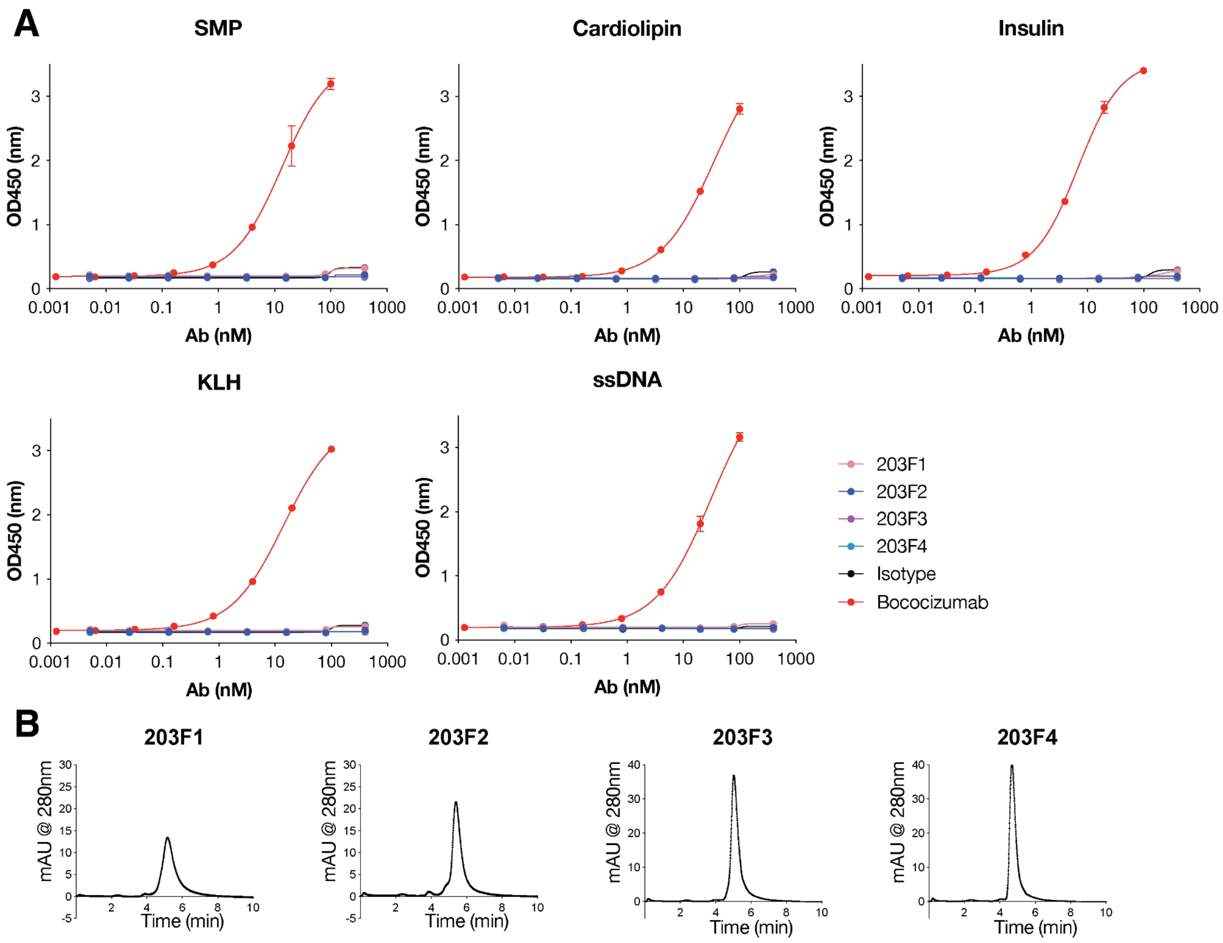


**Fig. S2**. **Polyspecific analysis of LRP8 Fabs. (A)** Polyspecific reagent (PSR)(1) ELISA binding of recombinant LPR8 Fabs indicated minimal cross-reactivity to a panel of autoantigens, including solubilized membrane proteins (SMPs), cardiolipin, insulin, keyhole limpet hemocyanin (KLH), and single-stranded DNA (ssDNA). The polyreactive antibody bococizumab(2) was used as a positive control, while anti-SARS-CoV-2 receptor binding domain (RBD) antibody CC12.1(3) served as a negative control. Binding was detected with peroxidase-conjugated goat anti-human IgG (H+L) antibodies. Data are mean + SD of two biological experiments. **(B)** Analytical size exclusion chromatography showing that LRP8 Fabs were monodispersed with regular column retention times.

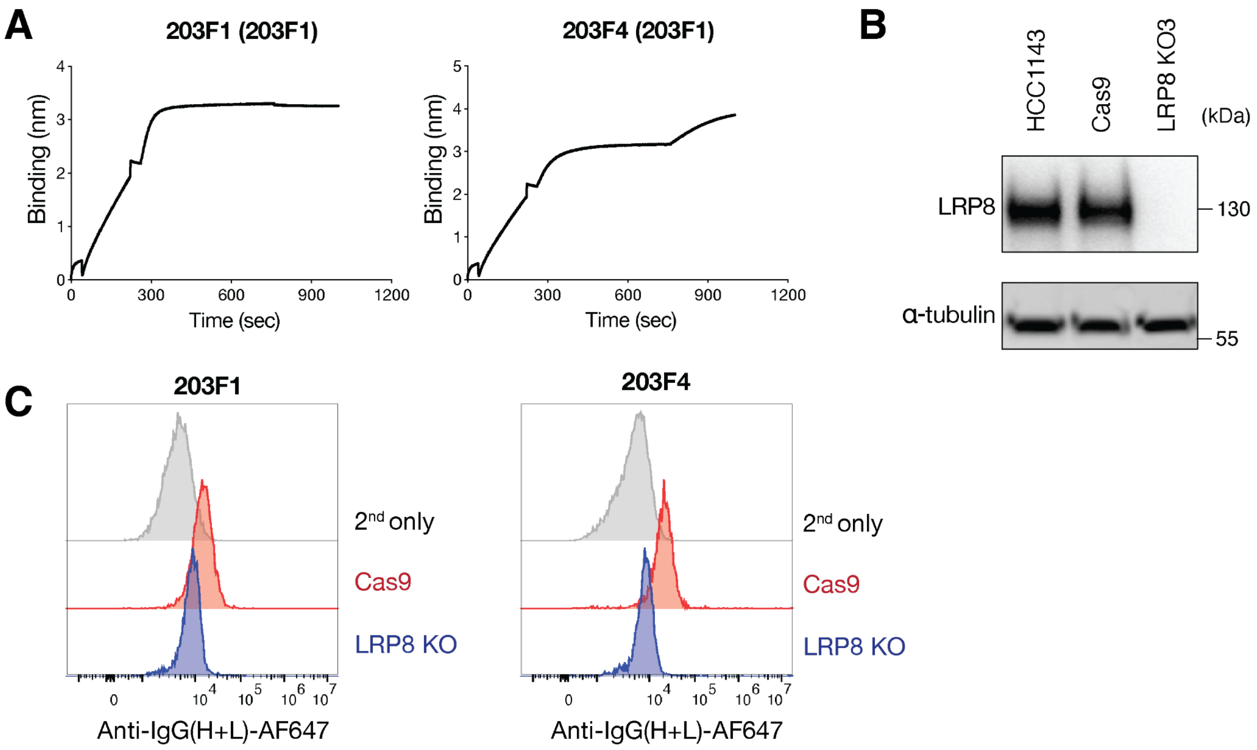


**Fig. S3**. **Epitope specificity and selectivity of LRP8 antibodies. (A)** Epitope binning revealed two different Fab binding epitopes on LRP8. Biotinylated LRP8β was captured using a streptavidin biosensor and a saturating concentration (200 nM) of 203F1 or 203F4 Fab were incubated for 10 min followed by incubation with 50 nM of the competing 203F1 Fabs for 5 min. **(B)** Western blot showed LRP8 knockout on HCC1143 cells. **(C)** Flow cytometry analysis revealed Fab binding to LRP8 positive HCC1143 sgRNA control (Cas9) cells but not LRP8 knockout (KO) cells(4). Cells were stained with LRP8 antibodies for 30 min, followed by AF647 conjugated anti-human IgG (H+L) antibody staining for 15 min. Data is representative from two independent experiments.

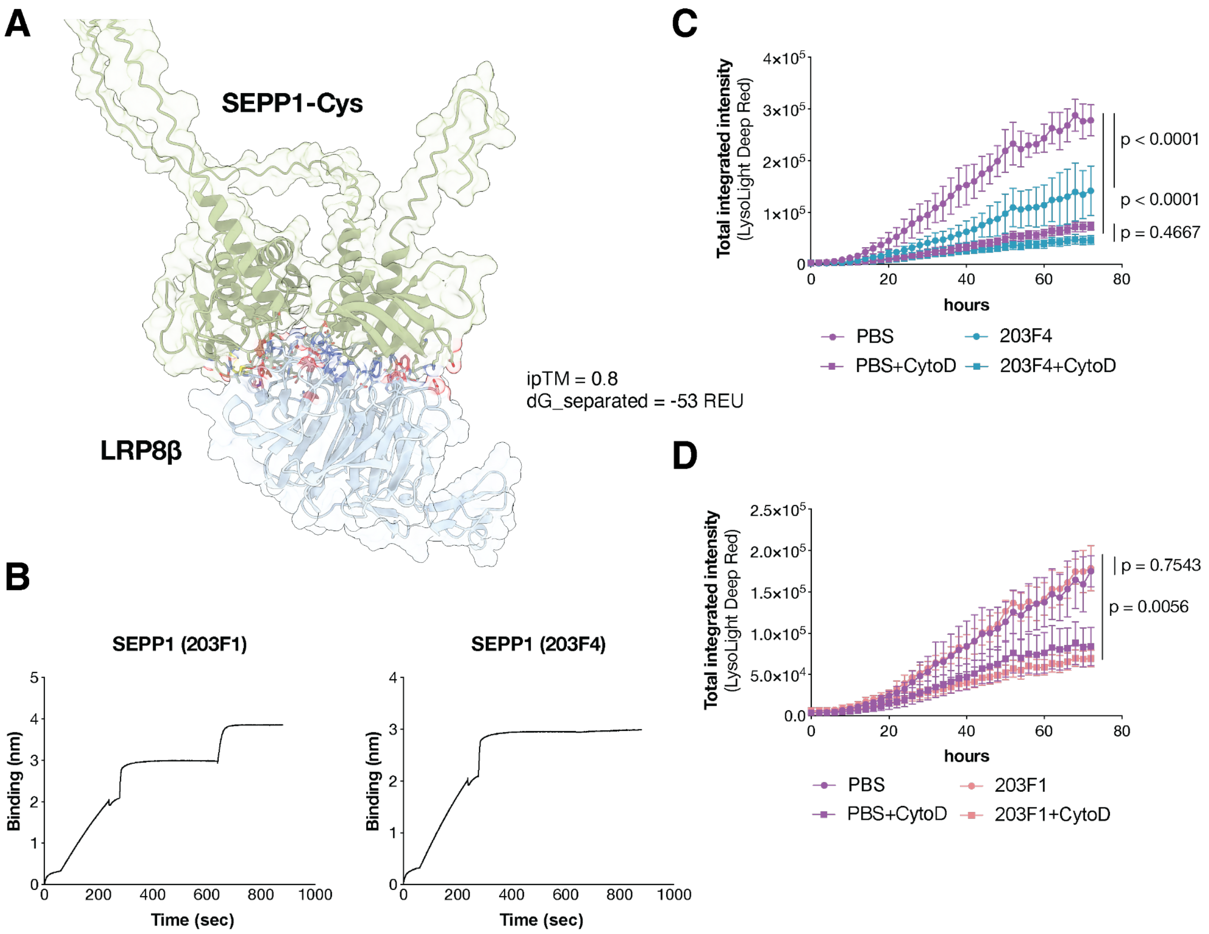


**Fig. S4**. **203F4 blocks selenoprotein P uptake.** **(A)** AlphaFold3 and Rosetta energy prediction of LRP8β interaction with SEPP1-Cys. Predicted complexes were visualized and analyzed using ChimeraX with a high-confidence interface (ipTM = 0.80). To evaluate binding energetics, AlphaFold-derived models were subjected to Rosetta-based scoring (dG_separated = -53 REU). **(B)** Epitope binning of SEPP1-Cys and LRP8 Fabs revealed two different epitopes on LRP8β. Biotinylated LRP8β was captured using a streptavidin biosensor and SEPP1-Cys at a concentration of 200 nM was incubated for 10 min followed by incubation with 50 nM of the competing LRP8 Fabs for 5 min. All incubation steps were performed in 1x PBS + 0.05% Tween + 0.2% BSA at room temperature. **(C-D)** Internalization of SEPP1-Cys in HCC1143 cells with or without macropinocytosis inhibition. SEPP1-Cys was labeled with the lysolight deep red dye (LLDR)(5) to track lysosomal trafficking. Cells were pre-incubated with 200 nM 203F4 Fab **(C)** or 203F1 Fab **(D)** for 30 min, in the presence or absence of 0.5 μM cytochalasin D (CytoD), followed by treatment with 50 nM SEPP1-Cys-LLDR. Images were captured every 2 h for 72 h on the Incucyte. Total integrated intensity was calculated by NIRCU x $\mu m^2$/image. Data represents mean ± SD from three biological replicates. Statistics were calculated by one-way ANOVA and Holm-Sidak multiple comparison tests.

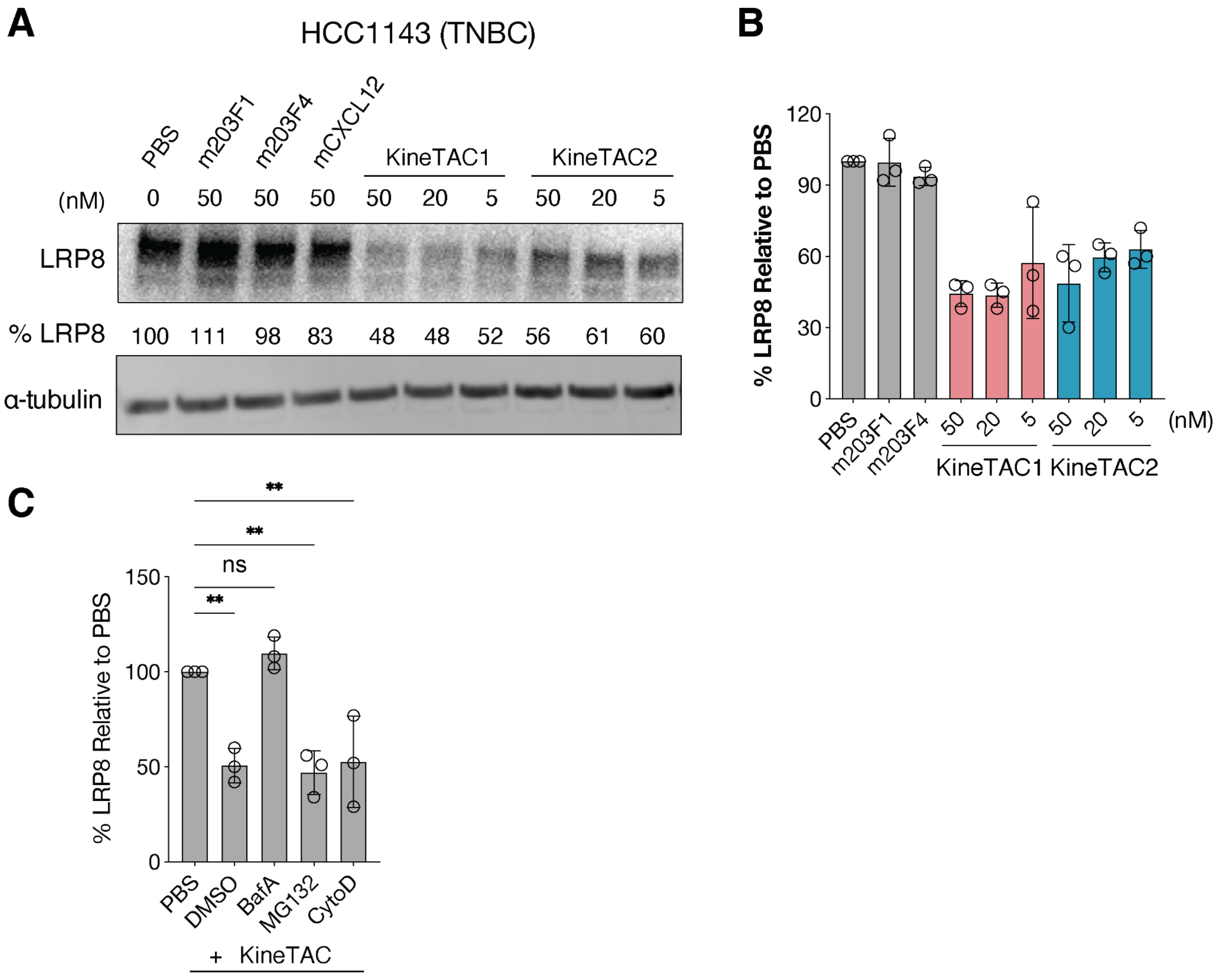


**Fig. S5**. **LRP8 degradation is lysosomal dependent. (A-B)** KineTAC-mediated LRP8 degradation in triple-negative breast cancer (TNBC) HCC1143 cells. Cells were treated with KineTAC for monomeric control antibodies (m203F1, m203F4) for 24 h. **(A)** Western blotting showed KineTAC-mediated degradation of LRP8 whereas monomeric antibody did not. **(B)** Percent LRP8 levels were quantified by ImageJ relative to the PBS control. Data represents mean ± SD from three biological replicates. **(C)** Cells were pretreated with either 500 nM Bafilomycin A (BafA), 500 nM MG132, or 500 nM Cytochalasin D (CytoD) for 1 h followed by 24 h treatment with 50 nM KineTAC. Percent LRP8 levels were quantified by ImageJ relative to the PBS control. Error bars represented standard deviations for three biological replicates. Statistics were calculated by one-way ANOVA. $^{**}P < 0.01$. ns, not significant.

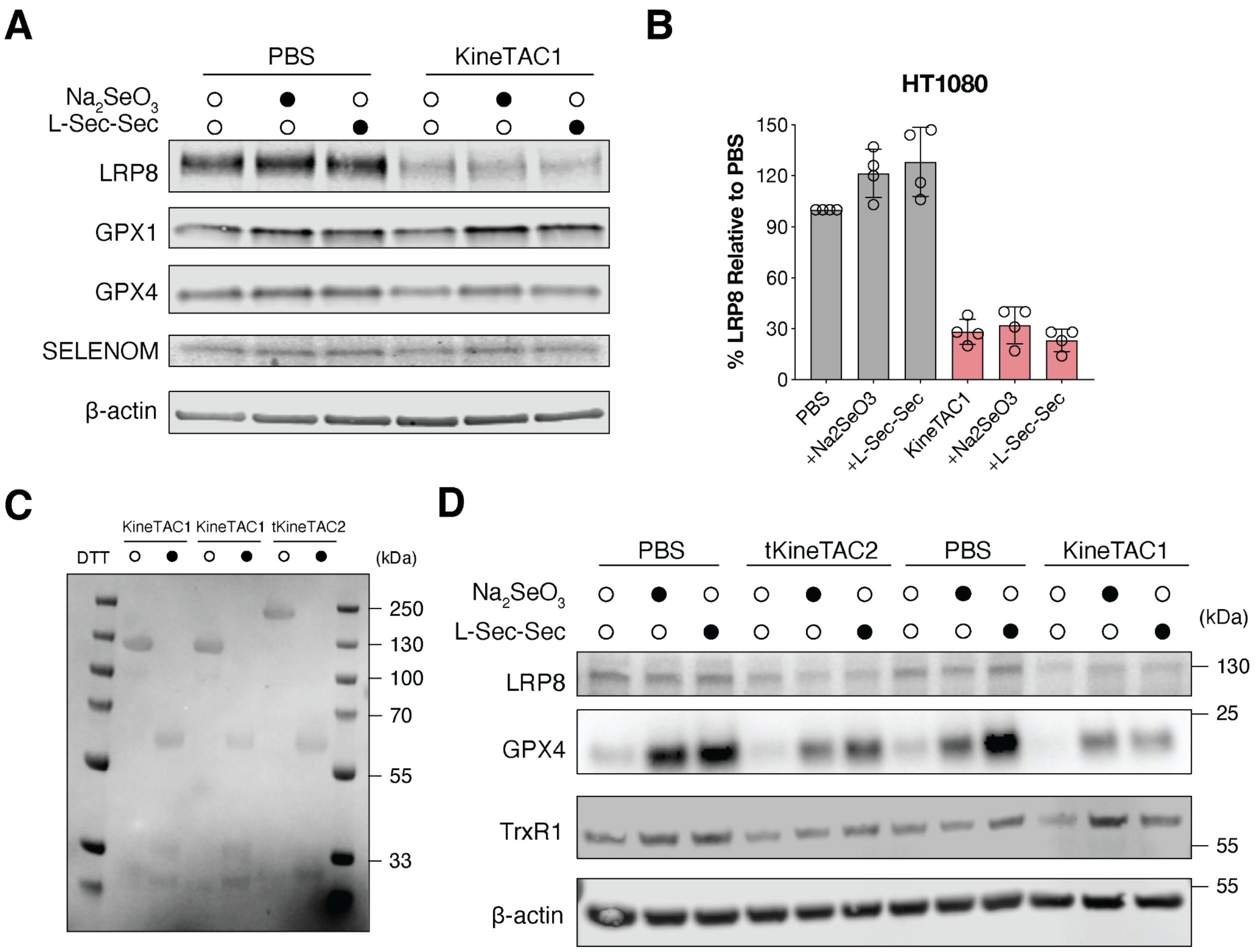


**Fig. S6**. **LRP8 degradation reduces selenoprotein levels. (A-B)** Western blotting analysis of HT-1080 cells treated with KineTAC1 in the presence or absence of selenium supplementation. 50 nM KineTAC1 treatment for 72 h efficiently degraded LRP8. Sodium selenite or L-selenocystine treatment didn't affect LRP8 degradation but rescued selenoprotein abundance. Percent LRP8 levels were quantified by ImageJ relative to the PBS control. Data represents mean ± SD from four biological replicates. **(C)** SDS-PAGE analysis of recombinant KineTAC1, KineTAC2, and tKineTAC2. Antibody samples were reduced with dithiothreitol (DTT) and denatured by boiling at 95 °C for 5 min. **(D)** Western blotting analysis of Kelly cells treated with KineTAC1 or tKineTAC2 for 72 h in the presence or absence of selenium supplementation. Whole cell lysates were blotted with LRP8, GPX4, TrxR1 and β-actin. Data are representative from at least three independent experiments.

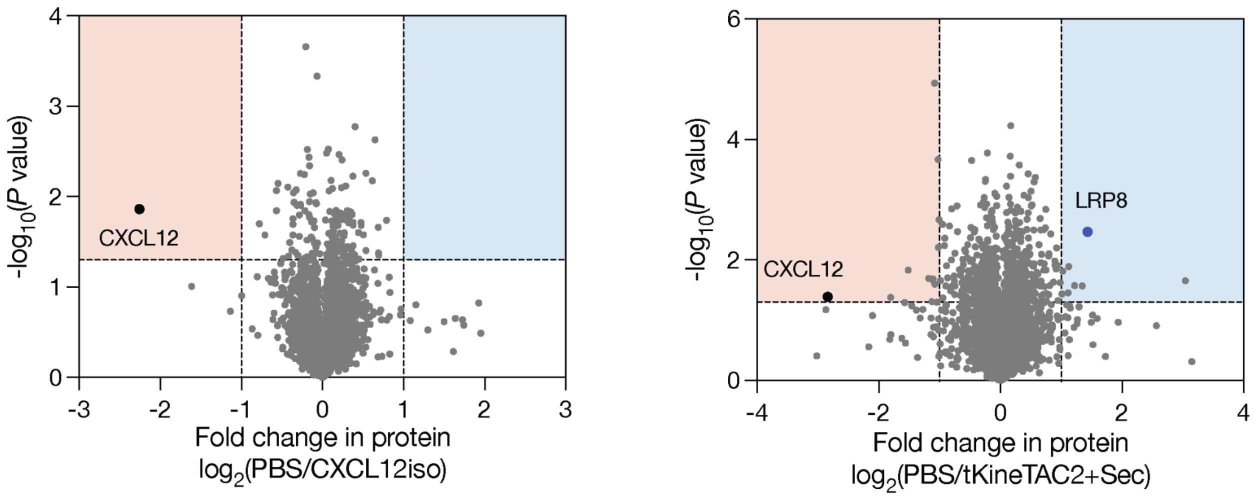


**Fig. S7**. **Whole cell proteomics analysis of Kelly cells following KineTAC treatment.** Kelly cells were treated for 48 hours with (left) 50 nM CXCL12 single-arm isotype antibody, or (right) 50 nM tKineTAC2 with 50 nM sodium selenite. Protein fold changes were quantified by mass spectrometry. Data are the mean of three biological replicates. P values were calculated using two-tailed Student's t-tests.

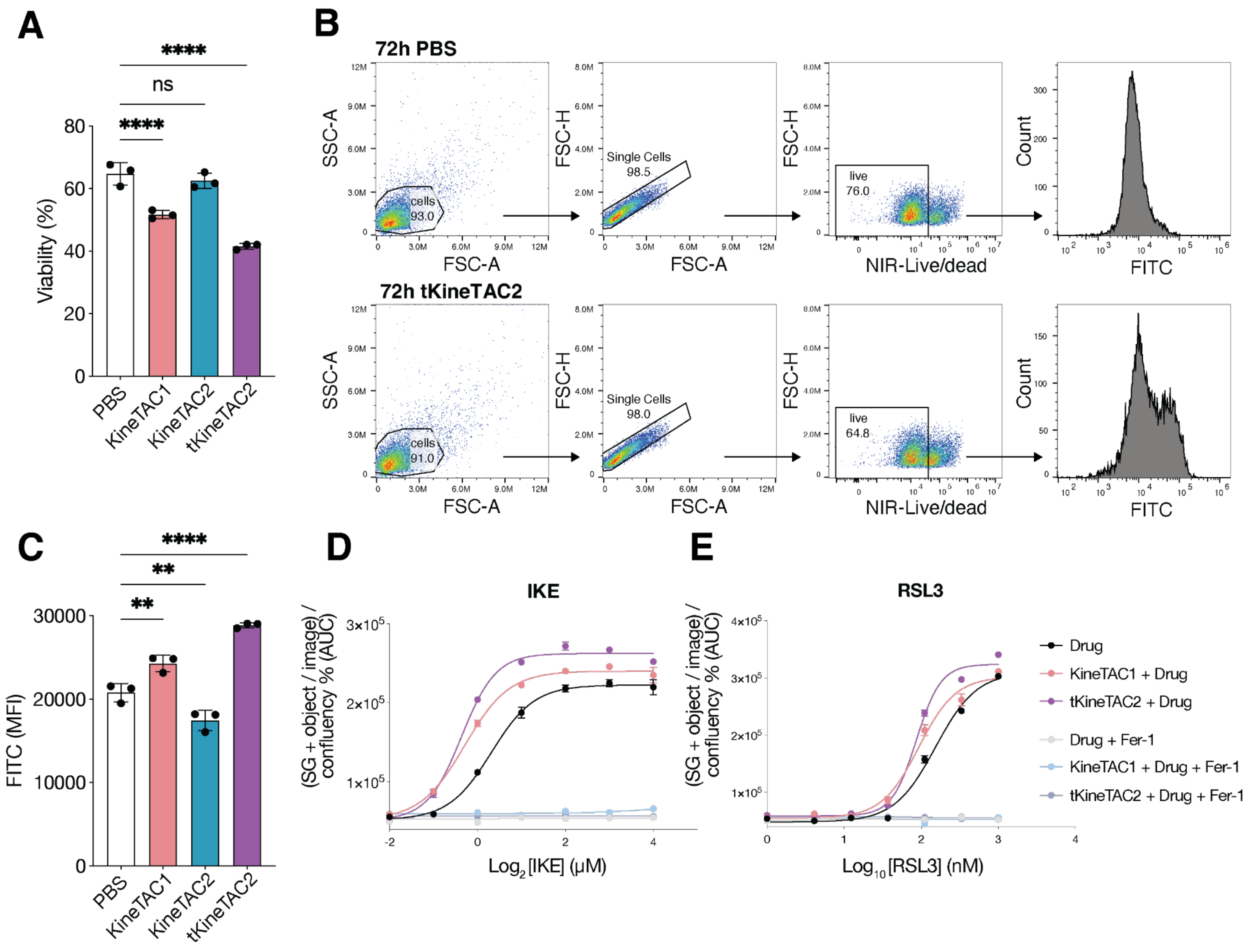


**Fig. S8**. **Targeting LRP8 increases tumor cell vulnerability to ferroptosis. (A-B)** Flow cytometry analysis of Kelly cell viability **(A)** and gating strategy **(B)** following KineTAC and RSL3 treatment. Kelly cells were pretreated with 50 nM KineTACs for 72 h, then incubated with 100 nM RSL3 for 5 h at 37 °C. Cells were stained with 5 μM BODIPY 581/591 C11 for 10 min at 37 °C, followed by Near-IR live/dead dye staining. Data represent mean ± SD of biological triplicates. **(C)** Flow cytometry analysis showing that KineTACs were sufficient to increase lipid peroxidation of Kelly cells without ferroptosis inducers. Cells were treated with PBS or 50 nM KineTACs for 72 h, followed by 5 μM BODIPY 581/591 C11 staining. Statistics were determined by one-way ANOVA. ***P* < 0.01. *****P* < 0.0001. ns, not significant. **(D-E)** Incucyte-based analysis of Kelly cells viability following 72 h treatment with KineTAC1 or tKineTAC2, in combination with IKE (**D**) or RSL3 (**E**), with or without Fer-1. Dead cells were labeled with the SYTOX green fluorescent dye. Data represent triplicate samples from three independent experiments.
